# HiCInterpolate: 4D Spatiotemporal Interpolation of Hi-C Data for Genome Architecture Analysis

**DOI:** 10.64898/2026.02.06.704438

**Authors:** H. M. A. Mohit Chowdhury, Oluwatosin Oluwadare

## Abstract

**Motivation:** Studying the three-dimensional (3D) structure of a genome, including chromatin loops and Topologically Associating Domains (TADs), is essential for understanding how the genome is organized, such as gene activation, cell development, protein-protein interaction, etc. Hi-C protocol enables us to study 3D genome structure and organization. Chromatin 3D structure changes dynamically over time, and modeling these continuous changes is crucial for downstream analysis in various domains such as disease diagnosis, vaccine development, etc. The high expense and impracticality of continuous genome sequencing, particularly what evolves between two timestamps, limit the most effective genomic analysis. It is crucial to develop a straightforward and cost-efficient method for constantly generating genomic data between two timestamps in order to address these constraints.

**Results:** In this study, we developed HiCInterpolate, a 4D spatiotemporal interpolation architecture that accepts two timestamp Hi-C contact matrices to interpolate intermediate Hi-C contact matrices at high resolution. HiCInterpolate predicts the intermediate Hi-C contact map using a deep learning-based flow predictor, and a feature encoder and decoder architecture similar to U-Net. In addition, HiCInterpolate supports downstream analysis of multiple 3D genomic features, including A/B compartments, chromatin loops, TADs, and 3D genome structure, through an integrated analysis pipeline. Across multiple evaluation metrics, including PSNR, SSIM, GenomeDISCO, HiCRep, and LPIPS, HiCInterpolate achieved consistently strong performance. Biological validation further demonstrated preservation of key chromatin organization features, such as chromatin loops, A/B compartments, and TADs. Together, these results indicate that HiCInterpolate provides a robust computer vision–based framework for high-resolution interpolation of intermediate Hi-C contact matrices and facilitates biologically meaningful downstream analyses.

**Availability:** HiCInterpolate is publicly available at https://github.com/OluwadareLab/HiCInterpolate.

## 1 Introduction

A chromosome is a DNA–protein complex composed of a long DNA molecule packaged with histone and non-histone proteins into chromatin, which carries genetic information within the nucleus of eukaryotic cells [1]. Chromosomes can interact with other chromosomes or with different loci on the same chromosome, giving rise to long-range chromatin contacts [2]. Together, these interactions organize the genome hierarchically in three-dimensional (3D) space. This 3D genomic organization plays a critical role in transcriptional regulation and encompasses several biologically significant features, including chromatin loops, A/B compartments, and topologically associating domains (TADs) [2, 3]. To study these 3D spatial features, chromosome conformation capture (3C)–based technologies were developed, with Hi-C, introduced by Lieberman-Aiden et al., becoming the most widely used protocol for measuring genome-wide chromatin interactions [2–4].

Recent studies have made important progress toward understanding the temporal regulation of 3D genome organization [5], yet capturing chromosomal dynamics across time at sufficient resolution and scale remains challenging [6–9]. According to Vian et al. [7], ATPases are necessary for the development of chromatin loops, which disappear after an hour in the absence of energy. When RAD21 dissolves away, these loops diminish, and they reappear three hours later when auxin is removed and gain strength over time [6]. Popay et al. recently demonstrated that NIPBL depletion within 24 hours results in the loss of RAD21, a cohesin core subunit, which in turn causes chromatin loops to disappear [9]. TADs become apparent at 2.5 hours after the cell emerges from the prometaphase arrest and during the cytokinesis stage. Over time, this region enhances interaction, and between 3 and 4 hours, long-range interactions denoted as A and B compartments become evident for several hours after entering into G1 [8]. To comprehend the underlying 3D spatial features discussed above, it is crucial to learn about cell cycle dynamics from interphase (G1, S, G2) to cytokinesis through the mitosis stage. This cell cycle and its underlying features are time-dependent, and sequencing this progression using the Hi-C protocol is expensive and complex.

Recently, 4DMax was developed to study dynamic chromosome conformation using time-series Hi-C data [10]. It reconstructs intermediate 3D chromosome structures by optimizing a likelihood function based on spatial restraints derived from contact matrices, producing smooth 4D models across time. However, this approach requires densely sampled time-series Hi-C data and involves computationally intensive optimization, limiting its scalability to high-resolution datasets. A related method, TADdyn, models structural rearrangements in time-series Hi-C data by reconstructing 3D chromatin structures from contact matrices available at each time point using molecular dynamics–based simulations [11]. However, it similarly depends on dense temporal measurements and computationally expensive simulations. More recently, HiCForecast [12] and HiC4D [13] applied video prediction frameworks to forecast future Hi-C data. However, these methods focus on temporal extrapolation rather than interpolation and do not explicitly reconstruct biologically consistent intermediate Hi-C states between sparsely sampled time points. Consequently, there is a pressing need for an efficient computational framework to interpolate high-resolution Hi-C data between two proximal time points, particularly in settings where dense time-series Hi-C experiments are infeasible, in order to improve the characterization of chromosomal dynamics.

In this study, we propose **HiCInterpolate**, a deep learning–based framework, inspired by [12, 14], which uses flow prediction between two input Hi-C maps and a U-Net–like architecture to interpolate high-resolution Hi-C data. Our approach achieved strong performance across multiple evaluation metrics, including SSIM, PSNR, GenomeDISCO, and LPIPS. We further demonstrate that the interpolated Hi-C data faithfully recapitulate key 3D genome features, including A/B compartments, topologically associating domains (TADs), and chromatin loops, in agreement with the corresponding ground truth data. Moreover, HiCInterpolate provides an integrated analysis pipeline that enables downstream biological analyses of the interpolated Hi-C data. To the best of our knowledge, HiCInterpolate is the **first computer vision–based framework** that effectively interpolates biologically meaningful intermediate Hi-C states between two observed time points at high resolution.

## 2 Material and methods

HiCInterpolate predicts intermediate Hi-C frame, 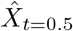 from two inputs, *X*_0_, *X*_1_, and provides an integrated pipeline for downstream biological analyses (Figure 1A). Our interpolation architecture is inspired by [14], where we implemented deep learning based bi-directional flow and UNet encoder and decoder to predict the finest and optimal frame (Figure 1B). Mathematically, we explain HiCInterpolate as (Equation 1),

**Fig. 1.**
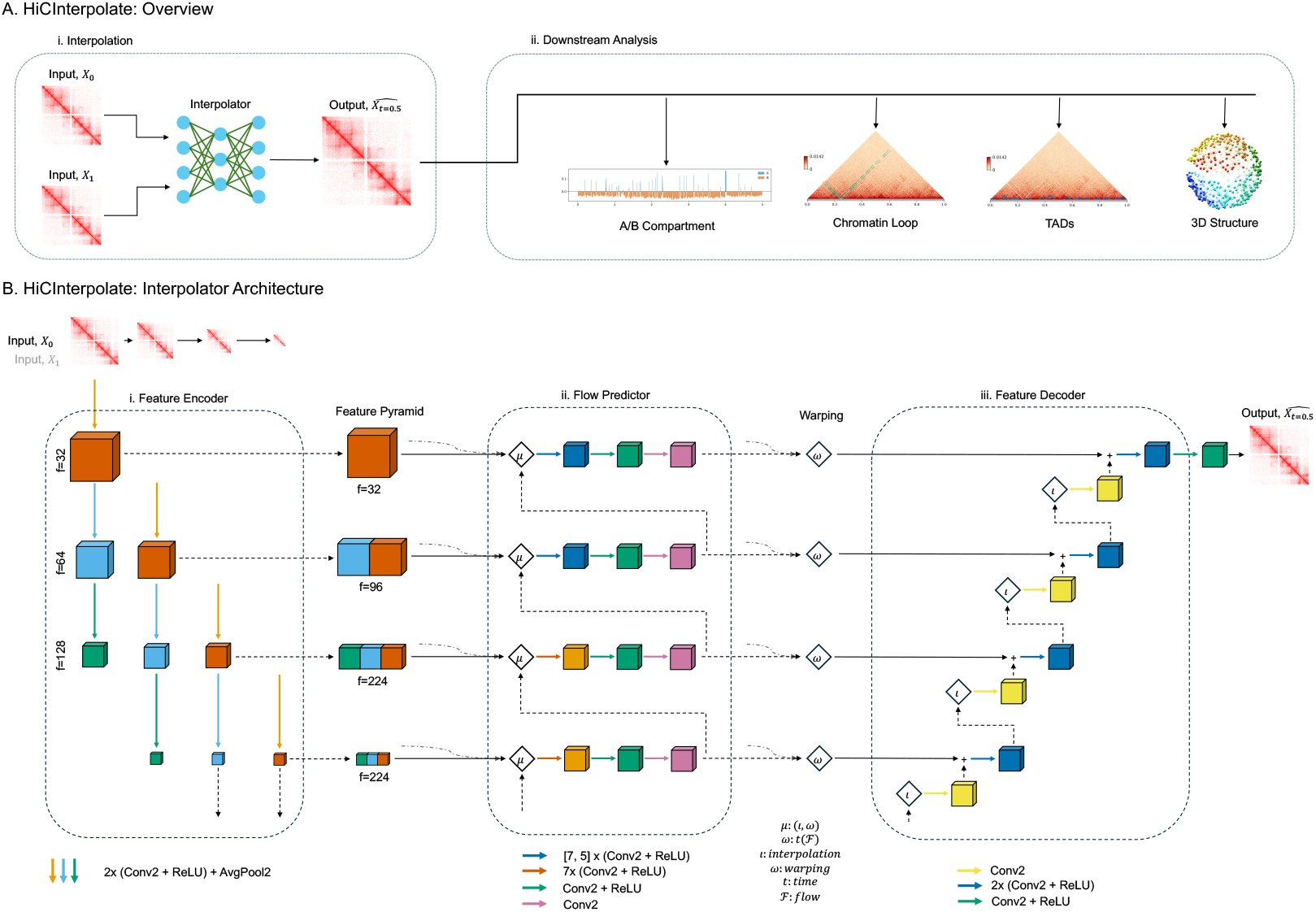
HiCInterpolate architecture. A. Overview of HiCInterpolate: i. Interpolation and ii. Downstream analysis, where it can perform: 1) A/B compartment analysis, 2) chromatin loop prediction, 3) TAD prediction, and 4) 3D structure prediction. B. Interpolator architecture explaining its different modules: i. Feature Encoder, ii. Flow Predictor, and iii. Feature Decoder, including Feature Pyramid and Warping.

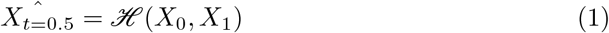

where ℋ = HiCInterpolate model architecture, *X*_0_ and *X*_1_ Hi-C contact matrix at timestamp 0 and 1 respectively, and 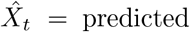 Hi-C contact matrix at timestamp, *t* = 0.5 (Figure 1A(i) - Interpolation). HiCInterpolate is infused with four different downstream analyses using the intermediate Hi-C contact matrix (Figure 1(ii) — Downstream Analysis). In the following subsections, we explain our model architecture’s three main modules—*i*. Feature Encoder, *ii*. Flow Predictor, and *iii*. Feature Decoder—in detail, along with dataset, data preprocessing, loss functions, and downstream analyses.

### 2.1 Dataset

In this study, we used various human datasets from different timestamps (Table 1). We used NIPBL-D7 cells with (i) DMSO and (ii) dTAG^*V*^ −1 treatment at different mitotic post-release timestamps [9]. HCT-116 is colorectal carcinoma cell line, and we used different timestamps of data with (i) auxin treatment (referred to as HCT-116), and (ii) RAD21-depleted and reintroduced (referred to as HCT116 - 2) [6]. Hela S3 (Replicate 1) cells are used at different timestamps during mitotic exit and G1 entry in prometaphase arrest condition [8].

**Table 1.**
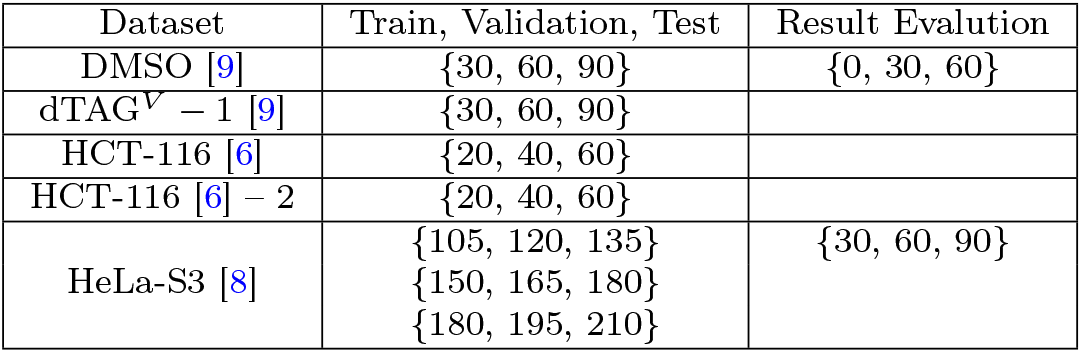
Dataset used to train, validate, test, and result evaluation. We used all chromosomes to train, validate, and test at 10Kb resolution. To evaluate the results, we used odd chromosomes from 11 to 21 at 10Kb resolution. The numeric values mentioned here are in minutes.

| Dataset | Train, Validation, Test | Result Evaluation |
| --- | --- | --- |
| DMSO [9] | {30, 60, 90} | {0, 30, 60} |
| dTAG<sup>V</sup> - 1 [9] | {30, 60, 90} |  |
| HCT-116 [6] | {20, 40, 60} |  |
| HCT-116 [6] - 2 | {20, 40, 60} |  |
| HeLa-S3 [8] | {105, 120, 135}<br>{150, 165, 180}<br>{180, 195, 210} | {30, 60, 90} |

### 2.2 Data preprocessing

We used different timestamps Hi-C datasets to train, validate, test, and evaluate our model (Data availability and Table 1). We converted *.hic* format data into a square symmetric contact matrix to feed into our model. First, we used the HiCExplorer [15] tool to convert *.hic* format data into *.cool* format at 10Kb resolution. During this conversion, we applied KR normalization to remove biases. Next, we used cooler [16] tool to extract *n* × *n* contact matrix from *.cool* format. After that, we applied Gaussian filter with *σ* = 1 to remove further noise for our model training, and *σ* = 1 ensures the spatial and biological features, such as loops, TADs, while averaging each bin contact with the nearest neighbor, improving the reproducibility [17]. Finally, we extracted patches of size 64 × 64, with a 20% overlap, from the *n* × *n* Hi-C contact matrix to serve as input to our model. This patch size was selected based on comparative experiments using 64 × 64, 128 × 128, 256 × 256, and 512 × 512 patches, where the 64 × 64 configuration achieved the best performance while avoiding computational overhead (Supplementary Data File 1).

### 2.3 Interpolator architecture

#### 2.3.1 Feature Encoder

HiCInterpolate takes two time-points input set, {*X*_0_}, {*X*_1_}, to predict the intermediate Hi-C frame, 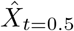. From the data preprocessing steps, HiCInterpolate accepts 64 × 64 patch size of a chromosome which we found to gives us optimal results (see Supplementary Data File 1). First, we create an input pyramid, 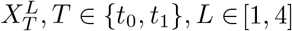, for the feature encoder. To create an input pyramid of four levels, we used 64 ∈ 64 input Hi-C image and gradually applied *AvgPool2d* with kernel size = 2 and *stride* = 2. Next, HiCInterpolate encodes features with a UNet encoder architecture inspired by [18]. In the feature encoder, we created a weight-shared feature pyramid for every input level, *L*, with a UNet encoder (Figure 1B(i) - Feature Encoder). From inputs 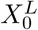 and 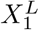, respectively, we derived features 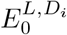 and 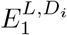, where *i* ∈ [1, 4] represents the depth of each level, *L* and *i* = 0 represents main input 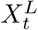. In terms of math, we stated as (Equation 2),

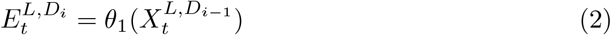

where *θ*_1_ represents the stack of two *Conv2D* and *ReLU* followed by *AvgPool2D*. In Feature Encoder, Figure 1B(i), every individual color (e.g. Vermillion = *D*_1_, Sky Blue = *D*_2_, Bluish green = *D*_3_) defines a different depth of convolution layers, and at the end of each convolution layer, we applied *AvgPool2D* to extract encoded features. At each depth, *D*_*i*_, we combined features from each level, *L*, to create multiscale features. At each level, *L*, encoded features share the same spatial dimension as other levels. At the end, we created a feature pyramid using multiscale features from each depth and level. We stated each feature map as (Equation 3),

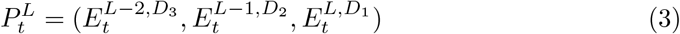

We constructed two feature pyramid for two input in Feature Encoder by stacked feature maps from each level preserving the same spatial dimension, where {Vermillion} at depth, *D*_1_, {Sky blue, Vermillion} at depth *D*_2_, {Bluish green, Sky blue, Vermillion} at depth *D*_3_, and so forth.

#### 2.3.2 Flow Predictor

After constructing the feature pyramid 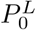 and 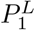 from two input, 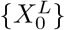 and 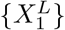 respectively, we estimate the forward residual flow, 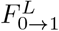 and backward residual flow, 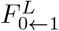 flow. To estimate residual bidirectional flow, we began at the coarse level inspired by [19]. First, we calculated the nearest interpolation using the previous level, *L*_*i*+1_, feature map from the two constructed feature pyramids, 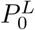 and 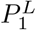. This interpolation up-samples the coarse feature map to align with the current feature map, as the coarse-level flow and finest-level flow will be the same [14], and we used the torch interpolation function. Then, we warped the upsampled flow with the current level, *L*_*i*_ feature map. Mathematically, we expressed it as (Equation 4),

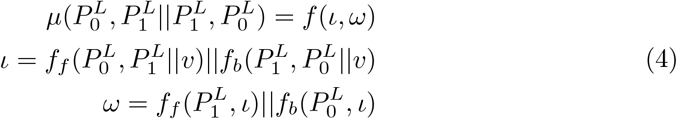

where *f*_*f*_ = forward flow, *f*_*b*_ = backward flow, and *v* = residue from the previous level. After calculating the flow, we applied a multi-convolution layer to predict the residual flow at each level. The top two stacked convolution layers are unique, and they do not share weights with other layers. The rest of the stacked convolution layer shares its weights with the predecessor layer. In our case, we stacked 7 (*Conv2D, ReLU*) for the shared weight depth, and for the top two unique depths, we stacked 7 and 5 (*Conv2D, ReLU*), respectively. In Flow Predictor, Figure 1B(ii)), the shared weight depth is indicated with Orange color, and the top two unique stack convolution layers are indicated with Blue color. After this layer, we constructed a *Conv2D, ReLU* layer (Blueish green) followed by another *Conv2D* layer (Redish purple) to predict the residual flow. Following this architecture, we constructed forward flow, 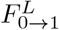, and backward flow, 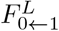. Mathematically, we express it as (Equation 5),

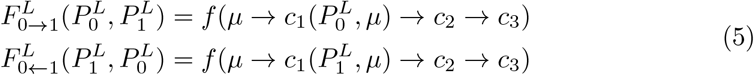

where *c*_1_ = {7, 5, 7} × (*Conv2D, ReLU*), *c*_2_ =(*Conv2D, ReLU*), and *c*_3_ =*Conv2D*. Finally, with this forward residual flow and backward residual flow, we synthesize the corresponding flow using nearest interpolation and apply time-dependent warping to predict the intermediate Hi-C contact matrix of each level. We express it as (Equation 6),

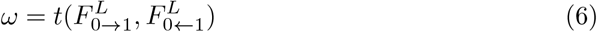

In our case, we multiplied both forward flow and backward flow by *t* = 0.5, concatenated them to predict the intermediate Hi-C contact matrix, and created an intermediate feature pyramid.

#### 2.3.3 Feature Decoder

Once we get the intermediate feature pyramid from the Flow Predictor step, we enhance the features from the coarse level to the finest level using a UNet decoder. First, we up-sample features from the previous level, applying nearest interpolation and pass these upsampled features to a *Conv2D* layer. After this, we concatenate these upsampled features and the current level feature, and finally pass this feature to a 2 (*Conv2D + ReLU*) layer to get the finest feature for this level. We expressed it as (Equation 7),

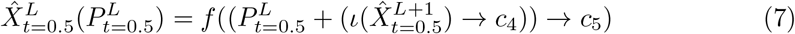

where 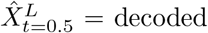 features, 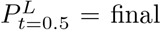 feature map from the Flow Predictor constructed pyramid, *c*_4_ = *Conv2D* layer and *c*_5_ = 2 (*Conv2D, ReLU*). Finally, we applied *(Conv2D, ReLU)* at the finest level to predict the final intermediate HiC contact matrix. In Feature Decoder, Figure 1B(iii), the Yellow color represents *Conv2D*, the Blue color represents 2 (*Conv2D, ReLU*) layers, and the Bluish green color represents (*Conv2D, ReLU*) layers.

#### 2.3.4 Loss Functions

In HiCInterpolate, we employed three different loss functions to supervise the predicted intermediate Hi-C contact matrix.

First, we used Mean Squared Erroe (MSE) loss function to minimize the pixelwise difference between the interpolated intermediate matrix, 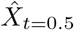, and the ground truth, *X*_*t*=0.5_. As HiCInterpolate predicts intermediate Hi-C data, we calculated MSE loss using a single channel and expressed it as (Equation 8),

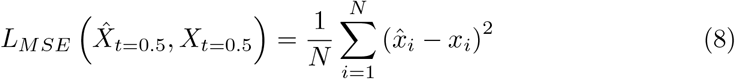

Second, we used Symmetry loss function to preserve the symmetrical structure. Hi-C contact matrix is a symmetrical diagonal matrix, and we expect each diagonal contact frequency to be the same. To preserve this feature, we calculated the symmetrical loss between the diagonal values on the predicted intermediate matrix, 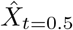, and stated it as (Equation 9),

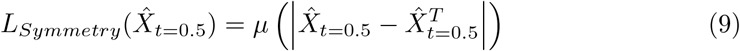

where *µ* = mean.

Finally, we used VGG loss, also known as perceptual loss, to improve the Hi-C intermediate matrix details [20]. We used a pretrained VGG19 with Imagenet_1k and employed layers (*l* ∈ {2, 7, 12, 21, 30}) to calculate the loss between the intermediate Hi-C matrix, 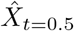, and the ground truth, *X*_*t*=0.5_, and stated as (Equation 10),

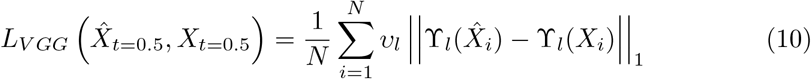

where ϒ_*l*_(*X*) ∈ ℝ^*H×W ×C*^ from each finest layer of the pretrained VGG19 model and *υ*_*l*_ = weight of every layers. As the Hi-C contact matrix has only one channel, we repeated this channel three times to make it compatible with the pretrained VGG19 model.

To extract the finest intermediate Hi-C matrix, we combined these three loss functions in our model with different weights for each loss function, and stated as (Equation 11),

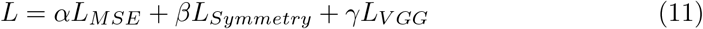

where *α* = weight for MSE loss, *β* = weight for Symmetry loss and *γ* = weight for VGG loss. This combined loss gives us an optimal result with a better evaluation metric curve.

### 2.4 Evaluation Metrics

To identify the best model from the training stages, we employed four evaluation metrics. (*i*.) The noise level between two Hi-C contact matrices is measured using Peak Signal-to-Noise Ratio (PSNR), which generates scores between 0 and ∞. The decibel (dB) unit is used, and a higher score is preferable.(*ii*.) Structural Similarity Index Measure (SSIM) [21] generates a score between −1 and 1 by comparing two Hi-C contact matrices while taking into account different fundamental image characteristics. (*iii*.) We also employed GenomeDISCO, a widely used evaluation metric in Hi-C analysis, which assesses the biological reproducibility between two Hi-C contact matrices and yields scores ranging from −1 to 1 [22].(*iv*.) HiCRep [23] is another biological reproducibility metric that computes the stratum-adjusted correlation coefficient (SCC) between two Hi-C contact matrices, with values also ranging from −1 to 1, and explicitly accounts for genomic distance.

During training, we used these metrics to validate our model and selected the optimal model weights based on their performance. In computer vision analysis, LPIPS [24] is a popular method for measuring perceptual image patch similarity based on pretrained deep neural networks. During the study of the results, we assessed the LPIPS score, which ranges from 0 (identical) to large (different).

### 2.5 Model Training, Validation, Testing and Evaluation Setup

We trained our model using the datasets listed in Table 1 under the *Train, Validation, Test* column. We generated single-channel 64 × 64 image patches to train the model. The 64 × 64 patch size was selected based on comparative experiments using 64 × 64, 128 × 128, 256 × 256, and 512 × 512 patches, where the 64 × 64 configuration achieved the best performance (Supplementary Data File 1).

The dataset was divided into training, validation, and test sets using an 80%, 10%, and 10% split, respectively. <u>Standard testing was performed on held-out patches from the same time points used during training and validation to assess within-distribution performance. In addition, a separate result evaluation was conducted on held-out time points (Table 1, *Result Evaluation* column) using odd-numbered chromosomes to assess temporal generalization within each dataset.</u>

During training, validation and testing, 64 × 64 patches were extracted along the diagonal of the Hi-C contact matrix, as these regions are enriched for informative short-to mid-range chromatin interactions. In contrast, during result evaluation, the model was applied to full Hi-C contact matrices and was not restricted to diagonal regions.

Model training was performed using the Adam optimizer with an exponential decay learning-rate scheduler, employing a maximum learning rate of 10^−4^ and a minimum learning rate of 10^−6^, with decay steps every 100 epochs. We used a composite loss function with variable weights; the initial loss weights were updated after 100 epochs. In Equation 11, we set *α* = 1.0, *β* = {0.1, 0.5}, and *γ* = {0.01, 0.1}, where *β* and *γ* were updated after 100 epochs. We trained the model for a total of 300 epochs with a batch size of 64 and selected the best-performing model based on validation metrics. The selected model, obtained at the 92nd epoch (Figure 2), achieved the following test-set performance: PSNR = 37.6263, SSIM = 0.9211, GenomeDISCO = 0.8014, HiCRep = 0.2185, and LPIPS = 0.1533.

**Fig. 2.**
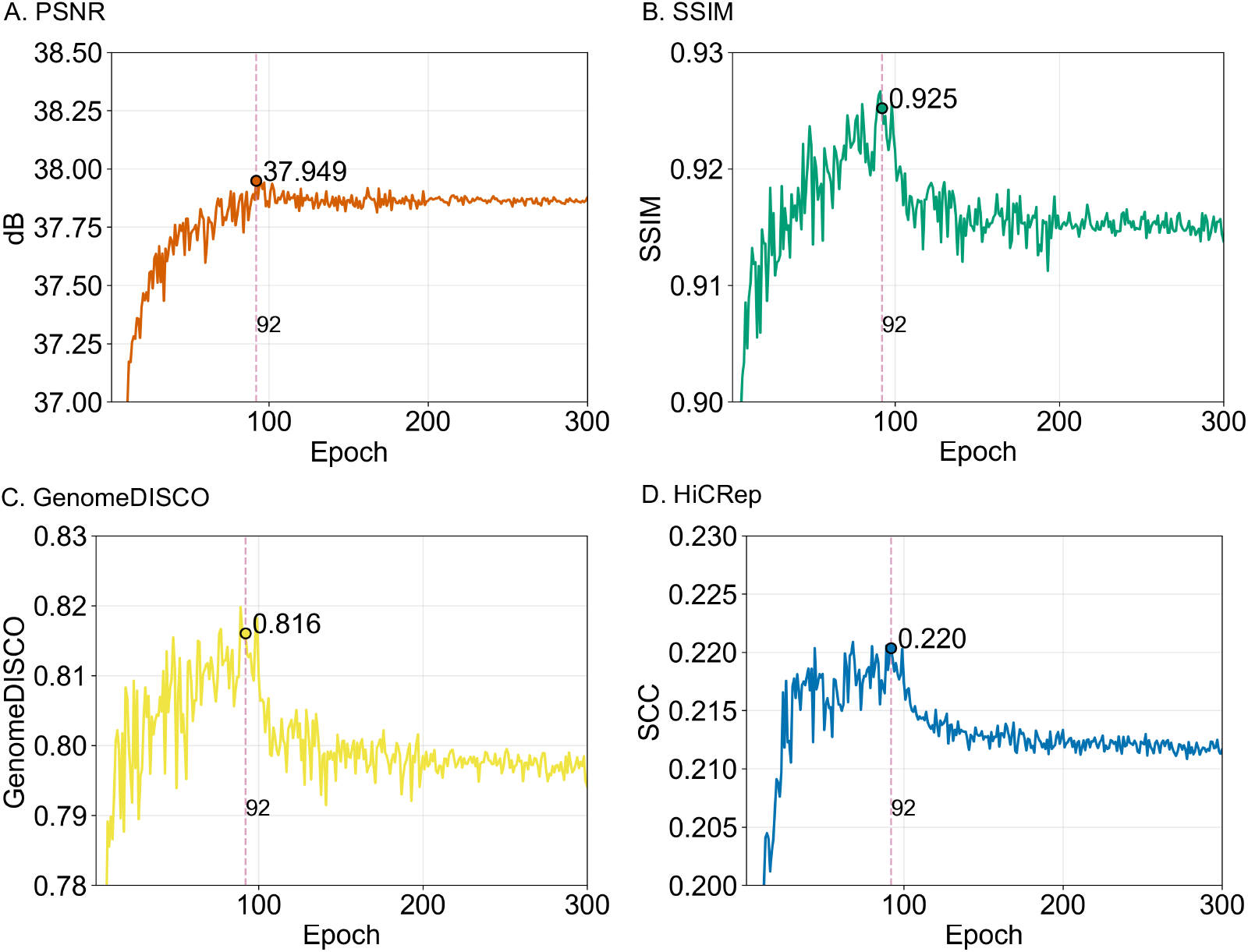
Validation scores during model training. During HiCInterpolate training, we performed 300 epochs and recorded the following scores: A. PSNR, B. SSIM, C. GenomeDISCO, and D. HiCRep, and at 92th epoch, we achieved an optimal score.

We used a dedicated workstation equipped with two NVIDIA RTX 4090 GPUs (24 GB VRAM each), 125 GB of system memory, and a multi-core CPU, running Ubuntu 24.04.3 LTS to train, test and validate our model.

### 2.6 HiCInterpolate Downstream Analysis

**We used the output of HiCInterpolate’s interpolator in conjunction with downstream analysis tools to provide four downstream biological analyses** Understanding the importance of downstream analysis of different genomic properties, we included A/B compartments, chromatin loops, TADs, and 3D structure prediction tools to make HiCInterpolation more robust and a useful tool for complete 3D genome analysis (Figure 1A - Downstream Analysis). In this analysis, we calculated the first principal component to differentiate between A and B compartments. Again, to predict chromatin loops, we incorporated HiCCUPS [4] in this analysis, as this is one of the most widely accepted tools for chromatin loop prediction. We included EmbedTAD [17] to predict topologically associating domain (TAD) regions, as it is a recently developed method based on graph embedding and clustering. Reconstruction of chromatin 3D structure is another important component of 3D genome analysis, as 3D structure visualization provides insights into chromatin organization and domain formation. To underscore this importance, we incorporated HiC-GNN [25] to predict chromatin 3D structure, as it is a recent deep learning–based method for chromosome 3D structure prediction.

## 3 Results

We assessed our trained model using DMSO dataset at *X*_0_ = 0 minutes, *X*_1_ = 60 minutes, and predicted 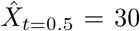 minutes and HeLa-S3 dataset at *X*_0_ = 30 minutes, *X*_1_ = 90 minutes, and predicted 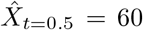 minutes Hi-C contact matrix at high resolution (Table 1 - Result Evaluation). In the Results section, we refer to **HeLa-S3** as a different dataset because the specific time points used for evaluation were not included during model training and are used to assess model generalization. In contrast, **DMSO** is referred to as a different timestamp evaluation, as the evaluated time points were only partially used during training. We used ground truth (DMSO: *X*_*t*=0.5_ = 30 minutes and HeLa-S3: *X*_*t*=0.5_ = 60 minutes) for our trained model assessment. It is worth noting that we could not compare against 4DMax [10] because it does not produce intermediate Hi-C contact matrices with consistent *n×n* dimensions, which are required for our interpolation-based evaluation.

We evaluated the effectiveness of our model using odd-numbered chromosomes (11, 13, 15, 17, 19, and 21). First, we assessed similarity and reproducibility using multiple metrics, including PSNR, SSIM, GenomeDISCO, HiCRep, and LPIPS. The resulting scores showed close agreement with those observed during training. Next, we performed biological validation of the model-generated Hi-C data. HiCInterpolate produced robust chromatin loops and topologically associating domains (TADs) that exhibited strong agreement with the ground truth. Finally, we visualized three-dimensional genome structures reconstructed from the model-generated Hi-C data, where HiCInterpolate consistently recovered structures highly similar to those derived from the original data. **Overall, our method generated high-quality intermediate Hi-C contact matrices while preserving key biological features (Figure 3A; Figure 4A)**.

**Fig. 3.**
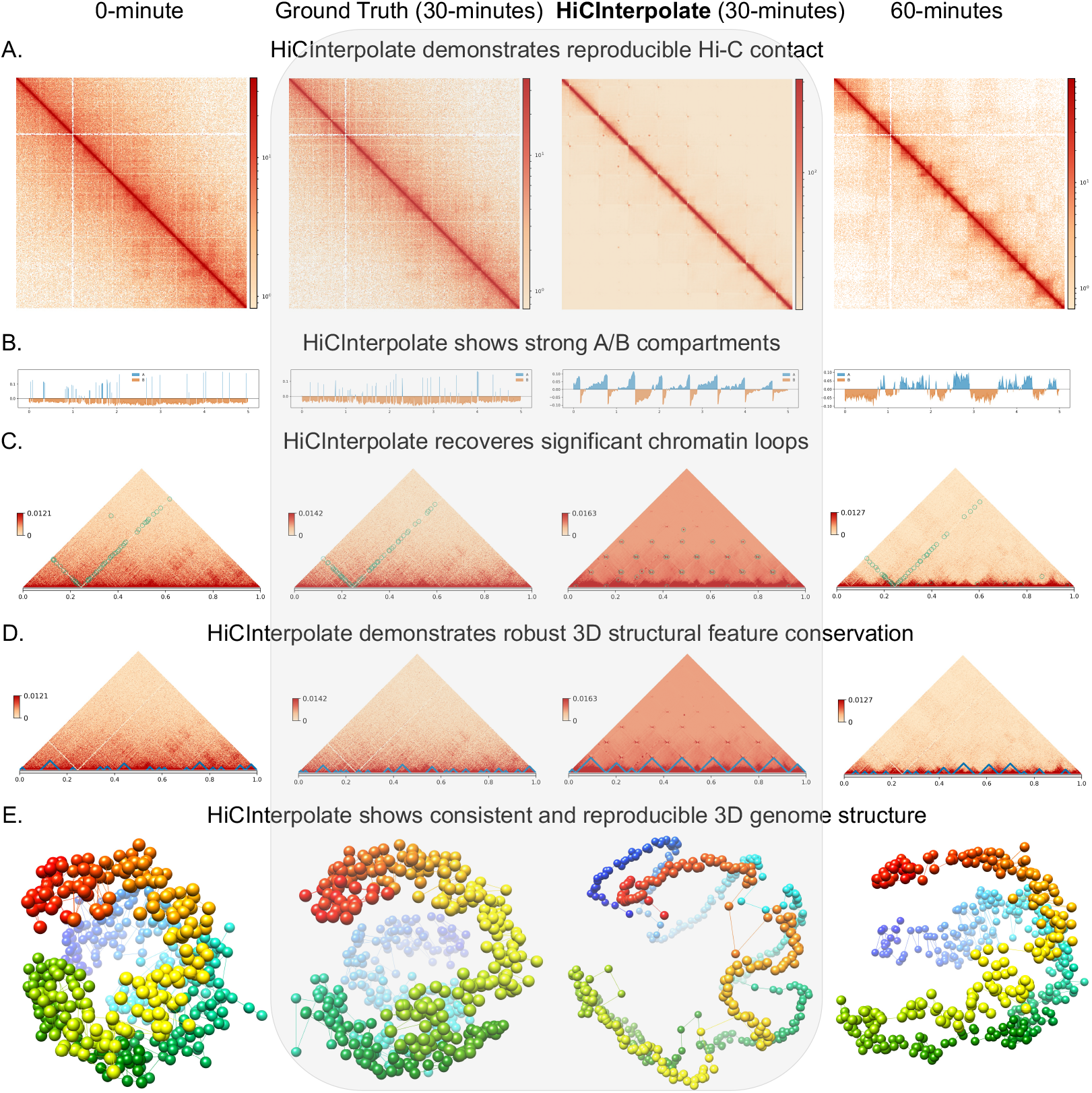
HiCInterpolate performance on different timestamps. A. HiCInterpolate predict intermediate Hi-C contact matrix of 30 minutes using DMSO chromosome 11 at 10Kb resolution. On the predicted Hi-C contact matrix, HiCInterpolate preserves B. A/B compartments, C. chromatin loops, D. TADs, and E. chromatin 3D structure from 8Mb to 8.5Mb region.

**Fig. 4.**
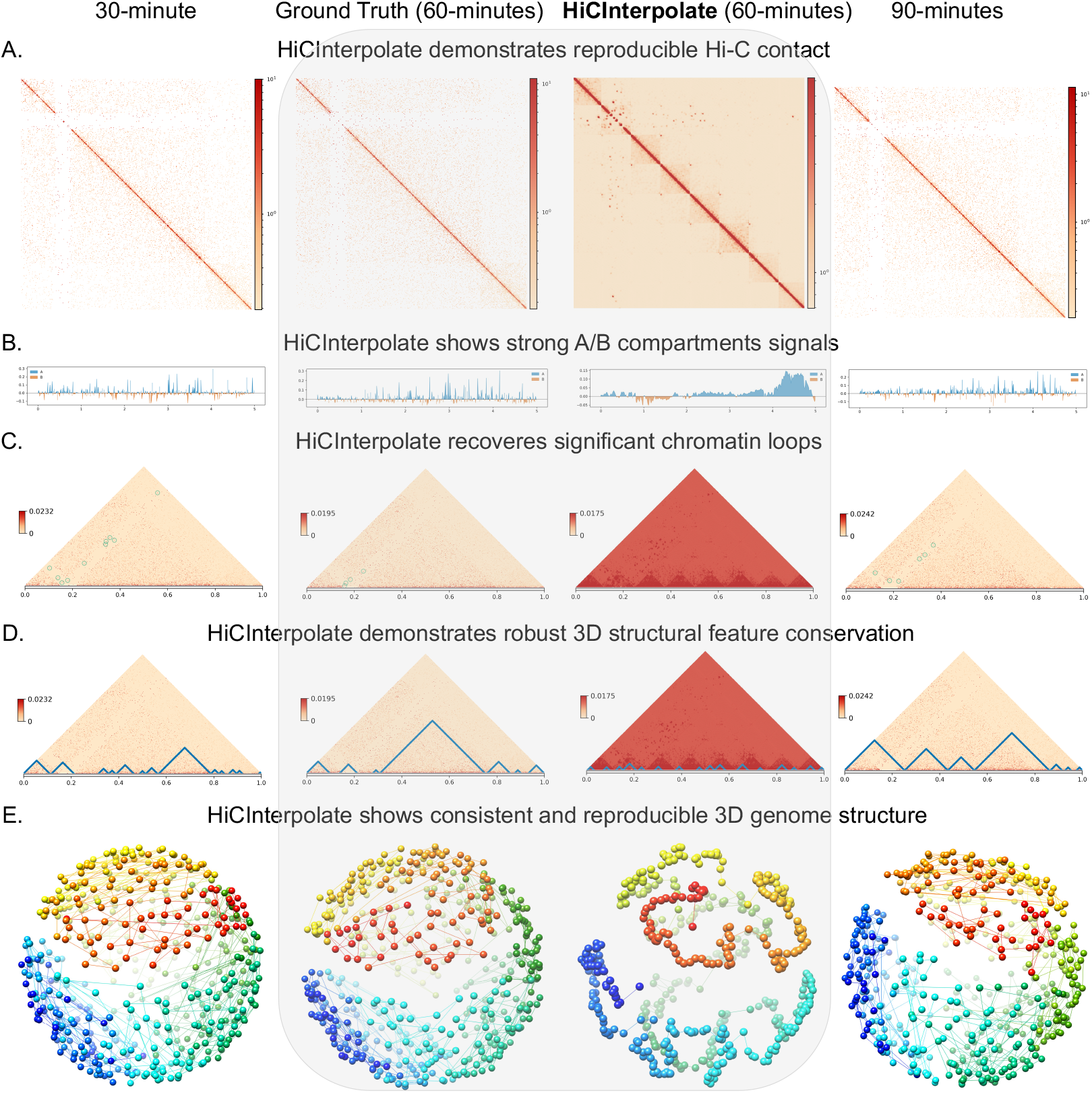
HiCInterpolate performance on different datasets. A. HiCInterpolate predict intermediate Hi-C contact matrix of 60 minutes using HeLa-S3 chromosome 11 at 10Kb resolution. On the predicted Hi-C contact matrix, HiCInterpolate preserves B. A/B compartments, C. chromatin loops, D. TADs, and E. Chromatin 3D structure from 8Mb to 8.5Mb region.

### 3.1 HiCInterpolate demonstrates high similarity and reproducibility

Using the two aforementioned datasets at 10 kb resolution, we evaluated HiCInterpolate’s predicted intermediate Hi-C matrices using PSNR, SSIM, GenomeDISCO, HiCRep, and LPIPS (Tables 2 and 3). PSNR and SSIM consistently achieved high values across chromosomes, indicating strong signal-level reconstruction quality and structural similarity to the corresponding ground truth across datasets and time points.

**Table 2.**
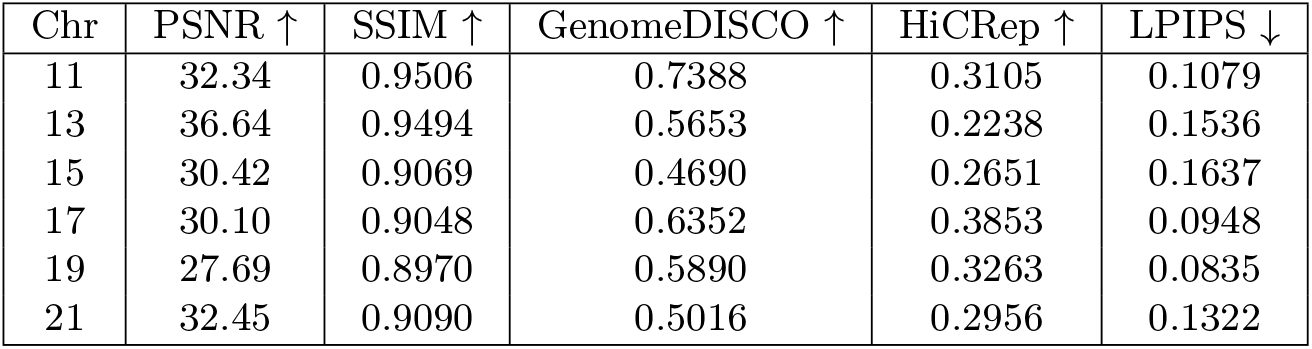
Similarity and reproducibility analysis on different timestamps. HiCInterpolate predicted Hi-C contact matrix of 60 minutes using DMSO at 10Kb resolution and showed consistently strong scores across 5 different evaluation metrics.

**Table 3.** Similarity and reproducibility analysis on different datasets. HiCInterpolate predicted Hi-C data of 60 minutes using HeLa-S3 at 10Kb resolution and showed consistently strong scores across 5 different evaluation metrics.

| Chr | PSNR ↑ | SSIM ↑ | GenomeDISCO ↑ | HiCRep ↑ | LPIPS ↓ |
| --- | --- | --- | --- | --- | --- |
| 11 | 40.61 | 0.9420 | 0.7191 | 0.0663 | 0.1335 |
| 13 | 38.66 | 0.9224 | 0.4654 | 0.0356 | 0.1679 |
| 15 | 38.33 | 0.8905 | 0.4463 | 0.0596 | 0.1904 |
| 17 | 40.01 | 0.9462 | 0.7532 | 0.1049 | 0.1424 |
| 19 | 31.89 | 0.8903 | 0.7676 | 0.0958 | 0.1227 |
| 21 | 37.17 | 0.9073 | 0.5049 | 0.0845 | 0.1599 |

The HeLa-S3 dataset (Table 3) exhibits higher PSNR and comparable SSIM values relative to the DMSO dataset (Table 2), consistent with smoother temporal transitions and reduced noise that facilitate accurate interpolation. These results further demonstrate that HiCInterpolate generalizes effectively to unseen temporal settings.

To assess biological reproducibility, we employed two complementary metrics to account for variance and to capture domain-level interaction patterns in Hi-C contact matrices. We evaluated model performance using GenomeDISCO, a widely accepted metric for measuring biological similarity between Hi-C datasets. Overall, HiCInterpolate achieved stable GenomeDISCO scores across chromosomes and datasets; however, we observed comparatively lower GenomeDISCO values for the HeLa-S3 dataset, reflecting increased biological variability at this time point rather than a loss of structural consistency. We further validated reproducibility using HiCRep, which explicitly accounts for genomic distance and domain structure. Similar to GenomeDISCO, HiCRep scores for HeLa-S3 were generally lower than those observed for the DMSO dataset, consistent with increased chromatin reorganization across time. Importantly, both GenomeDISCO and HiCRep exhibited coherent trends across chromosomes and time points, indicating that the interpolated Hi-C matrices preserve biologically meaningful chromatin organization despite increased temporal variability.

Lastly, we evaluated perceptual similarity using LPIPS, a deep learning–based metric. HiCInterpolate achieved low LPIPS scores, consistent with the validation and test results observed during training. Overall, HiCInterpolate demonstrates strong similarity and biological reproducibility across all evaluation metrics, regardless of chromosome, time point, or dataset.

### 3.2 HiCInterpolate shows strong A/B compartments and recovers significant chromatin loops

Chromatin contacts give rise to biologically significant structures such as chromatin loops, TADs, and A/B compartments [2–4]. The development of Hi-C has enabled the study of chromosome organization in three-dimensional (3D) space. To evaluate A/B compartment recovery, we computed the first principal component (PC1) from HiCInterpolate’s predicted intermediate Hi-C matrices. We observed that the interpolated Hi-C matrices exhibit clear A/B compartment patterns that are comparable to the ground truth across different time points and datasets, with compartment signals progressively converging toward future time points (Figures 3B and 4B). These regions also show strong PC1 signals, consistent with gene density and transcriptional activity [2].

We further evaluated HiCInterpolate by assessing chromatin loop recovery. Chromatin loops arise from intra-chromatin interactions constrained by CTCF extrusion barriers and play a key role in gene regulation [4, 26]. Using HiCCUPS [4] with an FDR threshold of *<* 0.001, we detected loops from both the interpolated Hi-C matrices and the ground truth. We validated DMSO predicted loops with RAD21 ChIP-seq data [9] and HeLa-S3 predicted loops with CTCF ChIP-seq data [27] to biologically quantify HiCInterpolate predicted Hi-C contact matrix. In Figure 5, we observed that majority of the loops predicted from the HiCInterpolate HiC contact matrix are supported by ChIP data.

**Fig. 5.**
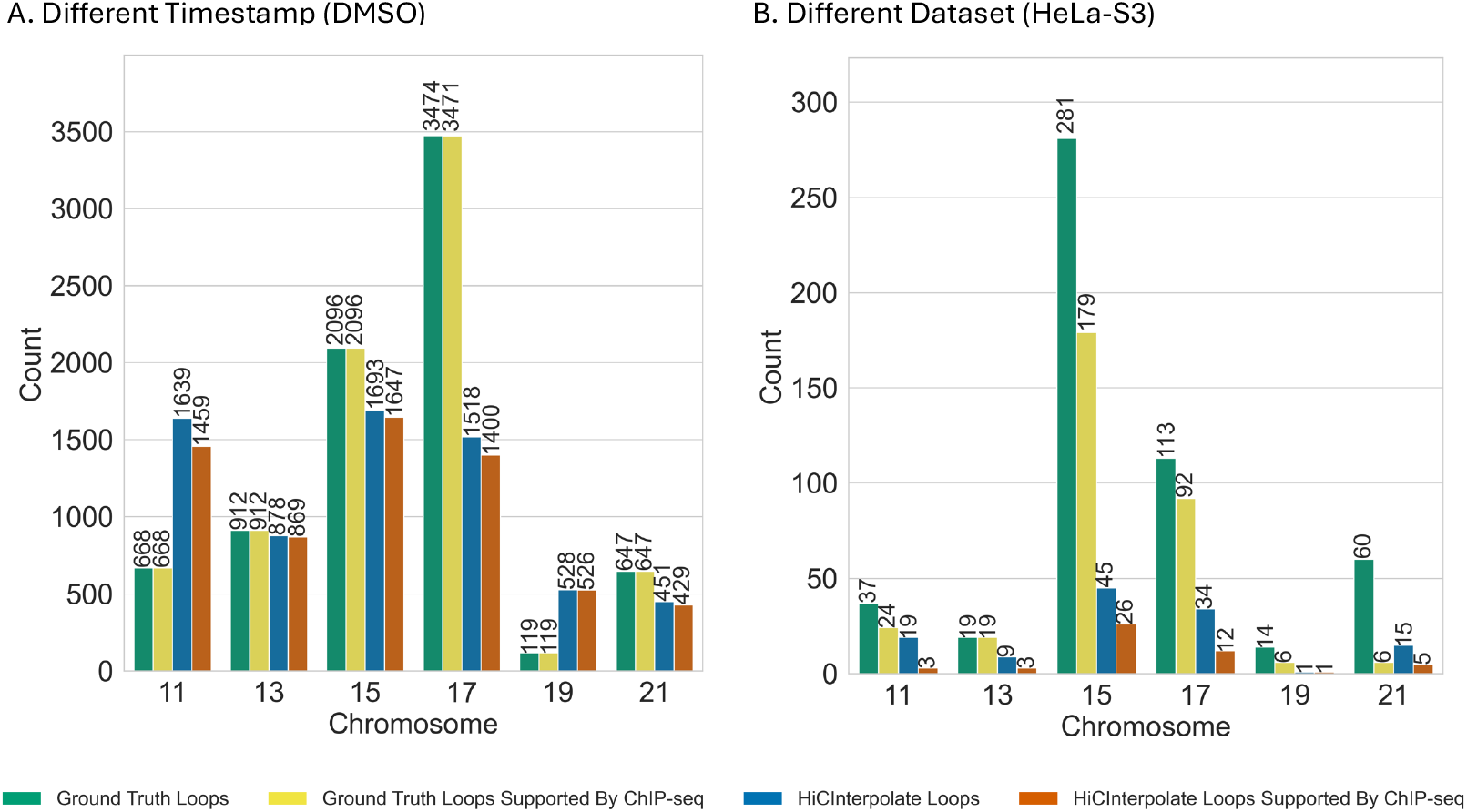
HiCInterpolate preserved chromatin loops across different timestamps and datasets. We validated predicted loops with RAD21 (for DMSO) and CTCF (for HeLa-S3) ChIPseq data and HiCInterpiolate preserved almost all biologically meaningful loops.

Overall, these results indicate that HiCInterpolate effectively recovers key chromatin organization features and provides biologically meaningful intermediate Hi-C matrices at high resolution.

### 3.3 HiCInterpolate demonstrates robust 3D structural feature conservation

Topologically associating domains (TADs) are self-interacting chromatin regions formed by intra-chromosomal interactions and play an important role in regulating gene expression [3]. To evaluate the ability of HiCInterpolate to preserve TAD structures in predicted intermediate Hi-C matrices, we applied three representative TAD detection tools—EmbedTAD [17], TopDom [28], and Spectral [29]—and compared the results with the ground truth.

We quantified TAD recovery using the Measure of Concordance (MoC) [30]. Figure 6 presents box plots of MoC score distributions across chromosomes, capturing variability across different TAD callers. While the distributions exhibit noticeable variability, HiCInterpolate maintains comparable median MoC ranges across chromosomes for both different time points (DMSO) and a different dataset (HeLa-S3), indicating consistent preservation of TAD structures. Absolute MoC values vary across tools, reflecting differences in TAD detection strategies; nevertheless, HiCInterpolate preserves the majority of ground-truth TAD structures (Figures 3D and 4D).

**Fig. 6.**
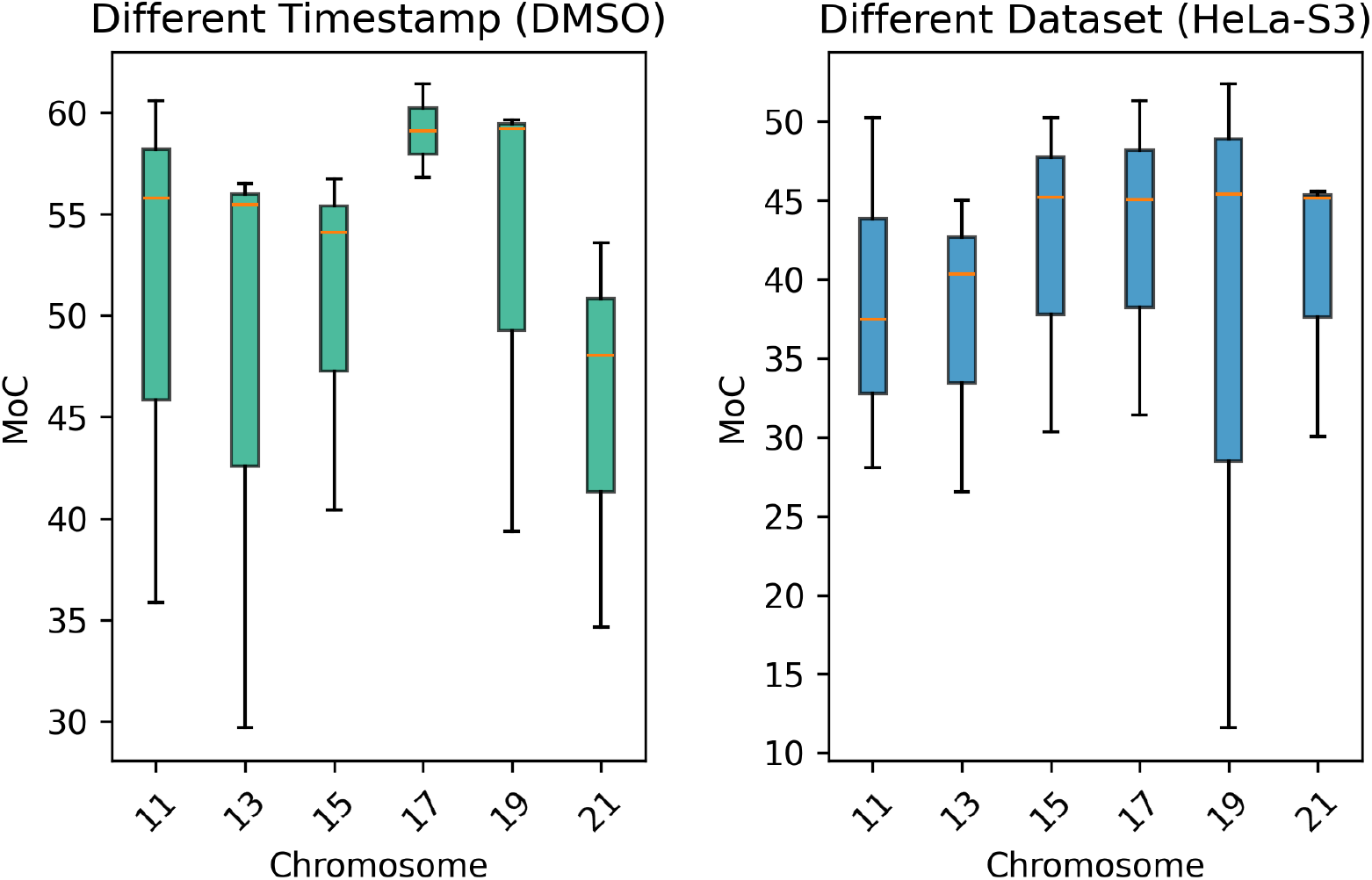
MoC score analysis from different TAD callers. Using 3 TAD callers, we computed the MoC score from different timestamps (DMSO) and different datasets (HeLa-S3) at 10Kb resolution. The box plots indicate that HiCInterpolate preserves TAD structures across chromosomes and experimental settings.

We further observed that MoC scores for the HeLa-S3 dataset were generally lower than those for the DMSO dataset, consistent with increased chromatin reorganization at different and unseen time points. Overall, these results indicate that HiCInterpolate conserves TAD-level chromatin organization across chromosomes, time points, and datasets.

### 3.4 HiCInterpolate shows consistent and reproducible 3D genome structure

Three-dimensional (3D) genome structure analysis helps us understand chromosomal features and their spatial organization, including TADs, chromatin loops, chromatin contacts, and protein-associated interactions, and the Hi-C protocol was developed to analyze chromosome 3D structure [4]. We visualized the 3D structures derived from HiCInterpolate’s interpolated intermediate Hi-C contact matrices to evaluate the performance of the proposed method. We utilized HiC-GNN [25] to predict and analyze 3D structures across different time points and datasets. HiC-GNN was trained separately on HiCInterpolate-generated Hi-C contact matrices and on the groundtruth Hi-C contact matrices, and 3D structures were predicted from each trained model. Using the predicted 3D coordinates, we visualized chromosome structures with Chimera [31] (Figures 3E and 4E). These visualizations demonstrate strong structural similarity between HiCInterpolate predictions and the ground truth.

To quantitatively validate 3D structural similarity, we computed Spearman’s correlation coefficient (SCC) between 3D structures reconstructed from HiCInterpolate-predicted intermediate Hi-C matrices and those derived from the ground truth. As shown in Figure 7, HiCInterpolate achieved consistently high SCC values for the DMSO dataset (SCC ≥ 0.59 across chromosomes), indicating strong preservation of 3D chromatin organization across different time points. For the HeLa-S3 dataset, SCC values remained moderate but stable (SCC ≥ 0.33), reflecting generalization to unseen temporal conditions and the increased reconstruction difficulty associated with sparser Hi-C contact maps. Overall, these results demonstrate that HiCInterpolate maintains robust 3D structural reproducibility while generalizing effectively across datasets.

**Fig. 7.**
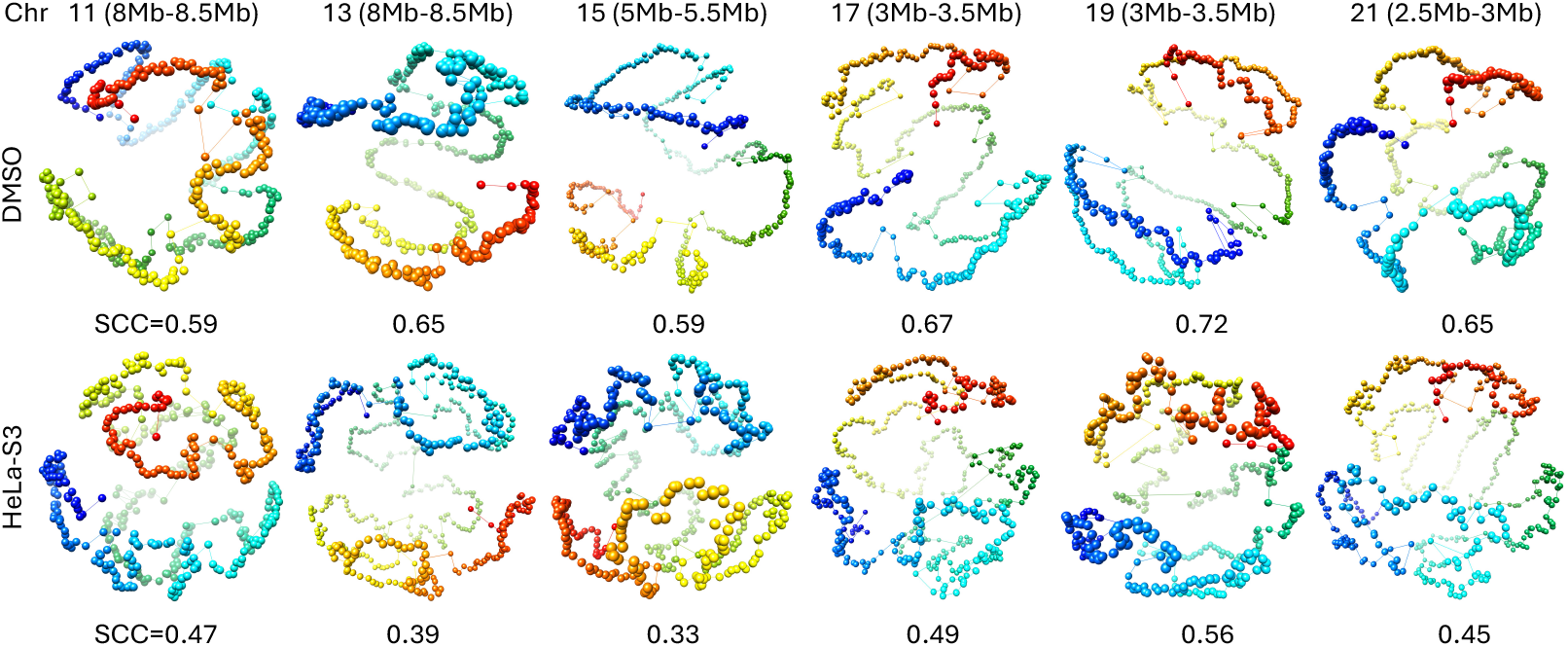
3D structure reproducibility analysis. We calculate the SCC score to measure the similarity between HiCInterpolate and the ground truth on different timestamps (DMSO) and different datasets (HeLa-S3) at 10Kb resolution. HiCInterpolate showed a significant similarity on DMSO and a considerable similarity on HeLa-3.

## 4 Discussion

In this study, we developed HiCInterpolate, a 4D spatiotemporal interpolation framework for predicting intermediate Hi-C contact matrices. HiCInterpolate employs a multi-level feature encoder, decoder, and flow prediction to interpolate high-resolution Hi-C matrices. We further integrated downstream 3D genome analysis tools for A/B compartments, chromatin loops, TADs, and 3D structure reconstruction, enabling comprehensive biological analysis of the interpolated Hi-C data.

HiCInterpolate demonstrated strong similarity, reproducibility, and biological consistency across multiple evaluation metrics, including PSNR, SSIM, GenomeDISCO, HiCRep, and LPIPS. Biological analyses showed high recovery of chromatin loops, robust A/B compartment signals, preservation of TAD structures as measured by MoC, and consistent 3D genome reconstruction as reflected by SCC scores. Overall, HiCInterpolate provides a robust computer vision–based framework for high-resolution interpolation of intermediate Hi-C contact matrices, enabling efficient and continuous downstream analysis between time points.

## 5 Data availability

We used in situ Hi-C human data for training, validation, and testing, as well as ChIP-seq (RAD21, CTCF) data. These data were downloaded from [4DN Data portal](https://data.4dnucleome.org/) and there accession numbers are GSE104334 [6], GSE133462 [8], and GSE277721 [9], GSE102884 [27]. All our processed Hi-C data patches used directly during training, validation, and testing are publicly available at https://doi.org/10.5281/zenodo.18303105.

## 6 Supplementary data

Supplementary Data File 1 contains validation metric scores during model training using (64 × 64), (128 × 128), (256 × 256), (512 × 512) diagonal patches from *n* × *n* Hi-C contact matrix.

## 8 Acknowledgments

Not Applicable.

## 9 Author contributions

H.M.A.M.C. conducted the analysis, wrote, and revised the manuscript, and O.O. conceived, wrote, revised the manuscript, and supervised this project. All authors reviewed the manuscript.

